# Whole-brain imaging and characterization of *Drosophila* brains based on one-, two-, and three-photon excitations

**DOI:** 10.1101/339531

**Authors:** Kuo-Jen Hsu, Yen-Yin Lin, Ann-Shyn Chiang, Shi-Wei Chu

## Abstract

To study functional connectome, optical microscopy provides the advantages of *in vivo* observation, molecular specificity, high-speed acquisition, and sub-micrometer spatial resolution. Now, the most complete single-neuron-based anatomical connectome is built upon *Drosophila*; thus it will be a milestone to achieve whole-brain observation with sub-cellular resolution in living *Drosophila.* Surprisingly, two-photon microscopy cannot penetrate through the 200-μm-thick brain, due to the extraordinarily strong aberration/scattering from tracheae. Here we achieve whole-*Drosophila*-brain observation by degassing the brain or by using three-photon microscopy at 1300-nm, while only the latter provides *in vivo* feasibility, reduced aberration/scattering and exceptional optical sectioning capability. Furthermore, by comparing one-photon (488-nm), two-photon (920-nm), and three-photon (1300-nm) excitations in the brain, we not only demonstrate first quantitative reduction of both scattering and aberration in trachea-filled tissues, but unravel that the contribution of aberration exceeds scattering at long wavelengths. Our work paves the way toward constructing functional connectome in a living *Drosophila*.

## Introduction

*Drosophila* is an important model animal to study connectomics since its brain is complex with 10^5^ neurons but still small enough to be completely mapped by optical microscopy with single-cell resolution. Compared to other model animals, the genetic tool box is more complete with *Drosophila*, and a connectome map based on *in vitro* structural registration of more than 30,000 cells has been established (Chiang et al., 2011), serving as an invaluable reference for functional connectome study. To study the functional connectome, two-photon fluorescence (2PF) microscopy is now the most popular tool because of its advantages on low photobleaching and phototoxicity, subcellular spatial resolution, and deep penetration depth (Helmchen & Denk, 2005). When observing living mouse or zebrafish brain with 2PF microscopy, the penetration depth approaches 1 mm, which is typically limited to about five scattering lengths (Helmchen & Denk, 2005; Horton et al., 2013). However, even using the same fluorophore and excitation wavelength, the reported imaging depths in a living *Drosophila* brain are much more limited. For example, when imaging GCaMP with excitation wavelength around 920-nm, activities from mushroom bodies (MB) had been recorded at only several tens of micrometers in depth (Honegger, Campbell, & Turner, 2011; Y. L. Wang et al., 2004). Using the same combination of laser and probe, the neuronal activities from antennal lobes (AL) are obtained with imaging depth less than 100 μm (Ignell et al., 2009; Root, Semmelhack, Wong, Flores, & Wang, 2007; Ruta et al., 2010; J. W. Wang, Wong, Flores, Vosshall, & Axel, 2003). Although the thickness of a *Drosophila* brain is only about 200 μm, which is much smaller than the typical imaging depth of 2PF microscopy in other model animals like mouse and zebrafish, to the best of our knowledge, no study has demonstrated *in vivo* whole-brain imaging in *Drosophila,* nor has characterized the image attenuation of a living *Drosophila* brain. The whole-brain observation capability is a major milestone toward establishing functional connectome in this model animal.

The underlying difficulty of living *Drosophila* whole-brain imaging is that, different from mouse and zebrafish, where blood vessels are responsible for oxygen exchange, air vessels, i.e., tracheae, are in charge of oxygen exchange in *Drosophila* brains. The micro-tracheae in the brain are a few micrometers in diameter (Beitel & Krasnow, 2000), comparable to near infrared wavelengths, and thus induce extraordinarily strong aberration/scattering from the air/tissue interface since the refractive index (RI) difference between air and tissue is much larger than that between blood and tissue. This tracheae-induced aberration/scattering impedes deep tissue observations inside a living *Drosophila* brain. However, the optical properties of trachea-filled tissues have not been well studied. Therefore, the aim of this work is to unravel the optical effect of tracheae, and to design a suitable method to increase imaging depth in trachea-filled tissues.

To increase imaging depth in living animals, there are several known approaches. For example, photo-activatable fluorophores (PAFs) have been used to suppress out-of-focus fluorescence (Wei, Chen, & Min, 2012), and high-energy lasers were used to enhance excitation efficiency at deep tissue (Theer, Hasan, & Denk, 2003). However, long converting time is required for PAFs, which is unfavorable to observe fast neural activities such as calcium dynamics, and high-energy lasers are potentially harmful due to multiphoton ionization. Adaptive optics (AO) is able to correct tissue-induced aberration, and has demonstrated significant image contrast and depth enhancement (Booth, 2014; Pedrazzani et al., 2016; Tang, Germain, & Cui, 2012; C. Wang et al., 2014). Nevertheless, the best depth achieved by AO in a living *Drosophila* brain to date is still less than 100 μm (Pedrazzani et al., 2016), since the aberration inside the insect’s brain is much larger than that of vertebrate’s brain (RI difference between air and tissue is at least ~ 1 order of magnitude larger than that between blood and tissue). In addition, typically AO does not compensate the scattering effect (Booth, 2014), which limits its impact in improving the imaging depth in *Drosophila*.

On the other hand, long excitation wavelength is well known to greatly improve penetration depth by substantially reducing scattering (Chu et al., 2003; Horton et al., 2013; Kobat, Horton, & Xu, 2011; Tao et al., 2017). In addition, the phase error caused by aberration (i.e., wavefront distortion) is inversely proportional to the excitation wavelength. The long wavelength approach is expected to reduce the amount of aberration caused by the RI difference, and thus enhancing imaging depth. Furthermore, high-order optical nonlinear excitations, such as three-photon absorption, are often combined with long wavelength, thus providing better excitation confinement, i.e. better optical sectioning capability, than 2PF, to improve image contrast in deep tissue. Combining these factors together, long wavelength excitation is promising for whole-brain imaging in *Drosophila*.

Here, two approaches are adopted to achieve whole-brain observation in a *Drosophila* brain. The first one is to pump our air inside the tracheae, i.e. degassing. Since the tracheae-induced aberration/scattering is largely removed in the degassed brain, 2PF microscopy penetrates through the whole brain. However, the *Drosophila* is not alive after degassing. To achieve *in vivo* whole-brain observation, the second approach is to use three-photon imaging based on excitation wavelength at 1300-nm in a GFP-labeled living *Drosophila* brain. The three-photon fluorescence (3PF) method provides exceptional excitation confinement and simultaneously reduced aberration/scattering, thus allows high-contrast and high-resolution image throughout the whole brain. The accompanying third harmonic generation (THG) modality provides detailed map of the densely distributed tracheae in the brain, useful for structure-function studies.

Furthermore, the optical attenuations of the *Drosophila* brains at various wavelengths are characterized, via 1PF (488-nm), 2PF (920-nm) and 3PF (1300-nm) excitation modalities. For the first time, the attenuations contributed from tracheae are quantitatively determined, with most contribution from aberration. We show that at short wavelength, scattering is the dominating attenuation; while at long wavelengths, the contribution of aberration can exceed that of scattering, and become the main impeding factor of whole-brain observation in *Drosophila* brains.

## Results and discussion

Within a living brain, the 1PF images lose the image contrast at around 40 μm (Video 1), as no structures are visible in the brain center where the white arrow points in the 60-μm panel of Figure 1A. The arrowheads indicate structures that located at the edge of the brain. To verify the effect of trachea, a brain is degassed, i.e. air in tracheae is pumped out. Figure 1B shows that 1PF in the degassed brain provides much better contrast in the center of brain at the same 60 μm depth, but cannot exceed 130 μm. The 1PF imaging depth of the degassed brain is comparable with that in mouse brains, which is mainly limited by scattering (Helmchen & Denk, 2005). Comparing the results in Figures 1A and 1B, it is obvious that degassing removes the additional attenuation contributed by tracheae.

**Figure 1.**
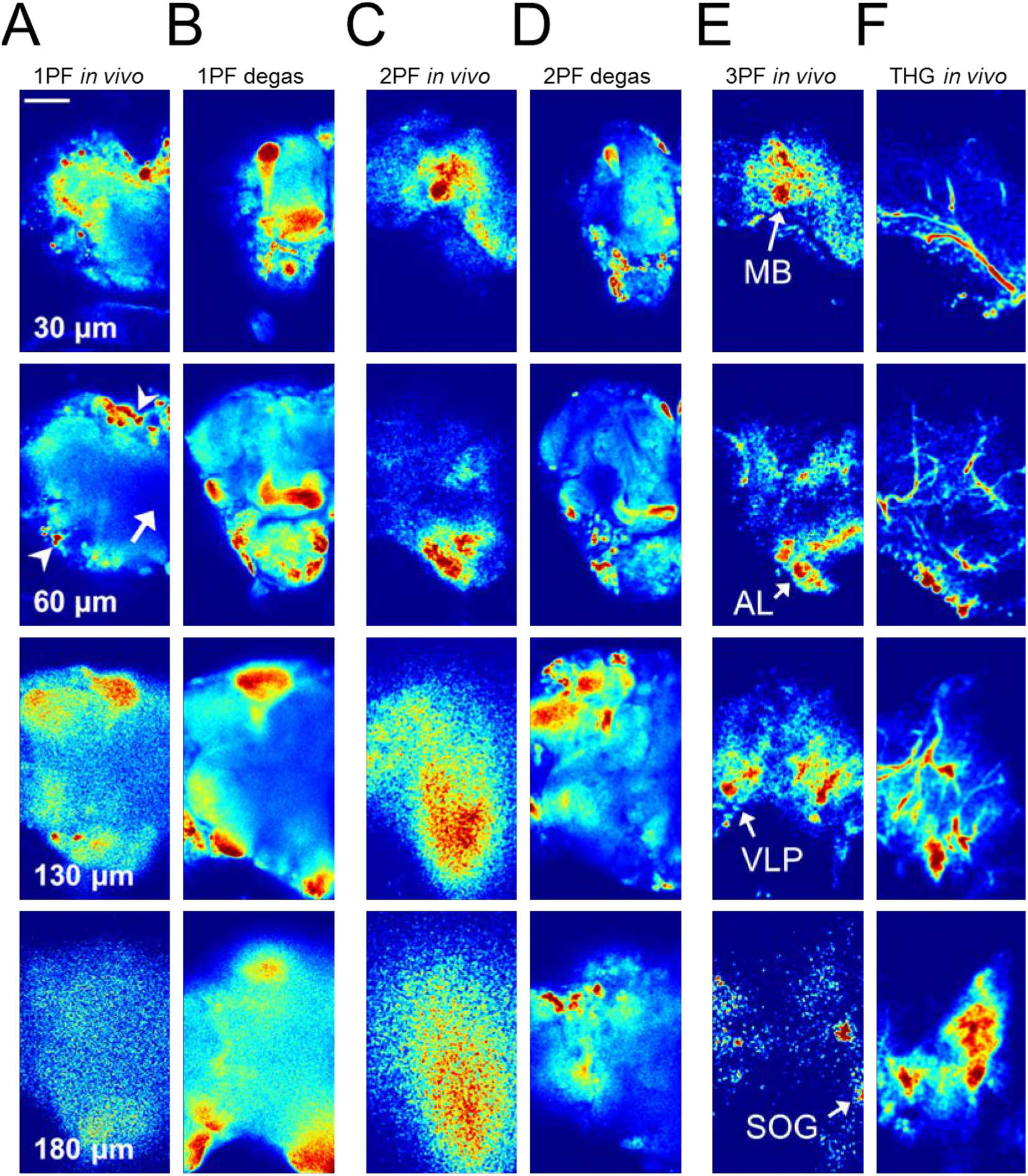
Living and degassed *Drosophila* brain images under different imaging modalities. (A), (C) and (E) are 1PF, 2PF and 3PF images at 30, 60, 130 and 180 μm depths of a living brain. (B) and (D) are 1PF and 2PF images of a degassed brain at the same depths. Although brain-edge structures (arrowheads) are still visible at 60 μm depth in (A), the image loses contrast in brain center (arrow). In the degassed brain, the penetration depth of 1PF is significantly improved in (B), but not approaching the bottom of brain. In (C), 2PF penetrates deeper than 1PF, but becomes blurry at depth beyond 100 μm. Whole-brain imaging is achieved in (D) by 2PF in a degassed brain, which nevertheless has no function. (E) shows only 3PF maintains contrast and reasonable resolution throughout the whole-brain *in vivo.* By comparing with structural connectome, LPUs can be identified in (E). MB: mushroom bodies; AL: antenna lobes; VLP: ventrolateral protocerebrum; SOG: subesophageal ganglion, which is the deepest LPU in a *Drosophila* brain. (F) THG images, a complementary contrast with 3PF, in the living brain show clear tracheae distribution. All the images are intensity normalized with individually brightest 1% pixels. Scale bar: 50 μm. Complete depth-images are given in Videos 1 - 3.

On the other hand, it is well known that using long excitation wavelengths with 2PF modality efficiently improves imaging depth, approaching 1 mm in mouse brains (Helmchen & Denk, 2005). Using the same excitation wavelength (~ 920-nm) and contrast agent (GFP families), Figure 1C shows the 2PF imaging depth in a living *Drosophila* brain indeed increases compared to Figure 1A, but reaches only about 100 μm, which is not adequate to penetrate the whole brain, mainly due to the tracheae.

By combining 2PF modality with a degassed brain, Figure 1D presents the first whole-brain imaging in a *Drosophila*. Reasonable contrast and resolution are maintained throughout the nearly 200 μm depth, manifesting that the trachea-induced aberration/scattering is the major restraint for deep-brain imaging in this model animal. However, please note that the animal is no longer alive after degassing. Moreover, the degassing process distorts the brain structures, making its association with the *Drosophila* structural connectome difficult.

In order to enable the study of whole-brain functional connectome, 3PF at excitation wavelength ~ 1300-nm was adopted. Figure 1E shows that this imaging modality provides subcellular resolution throughout a living *Drosophila* brain. Associating the 3PF images with an existing *Drosophila* brain database (*FlyCircuit,* http://www.flycircuit.tw), several local processing units (LPUs) (Chiang et al., 2011) are identified and marked by the white arrows.

During the three-photon excitation process, a complementary contrast, i.e. THG, is generated by the interfaces with RI differences, and thus is sensitive to the tissue/air boundaries in tracheae. Figure 1F shows that THG signals reveals the detailed distribution of tracheae, which are randomly distributed inside the brain and causing strong aberration/scattering.

To quantify signal attenuation coefficients (*μ_att_*) of different imaging modalities, Figures 2A - 2C show the decay of 1PF, 2PF, and 3PF signals, corresponding to Figures 1A - 1E respectively, within living or degassed brains (see Materials and Methods and Figure supplement 1 for derivation). Apparently, both degassing and long wavelength excitation help to reduce *μ_att_* significantly. The *μ_att_* of 2PF and 3PF are half and one-third of that of 1PF in living *Drosophila* brains, while the lowest *μ_att_* is provided by 2PF plus degassing. Comparing to mouse brain whose *μ_att_* of 2PF and 3PF are less than one-third and one-eighth of 1PF (Horton et al., 2013), the influence of longer excitation wavelength is much smaller in *Drosophila* brains. To better understand the results, the *μ_att_* has to be decomposed, as we explain in the following.

**Figure 2.**
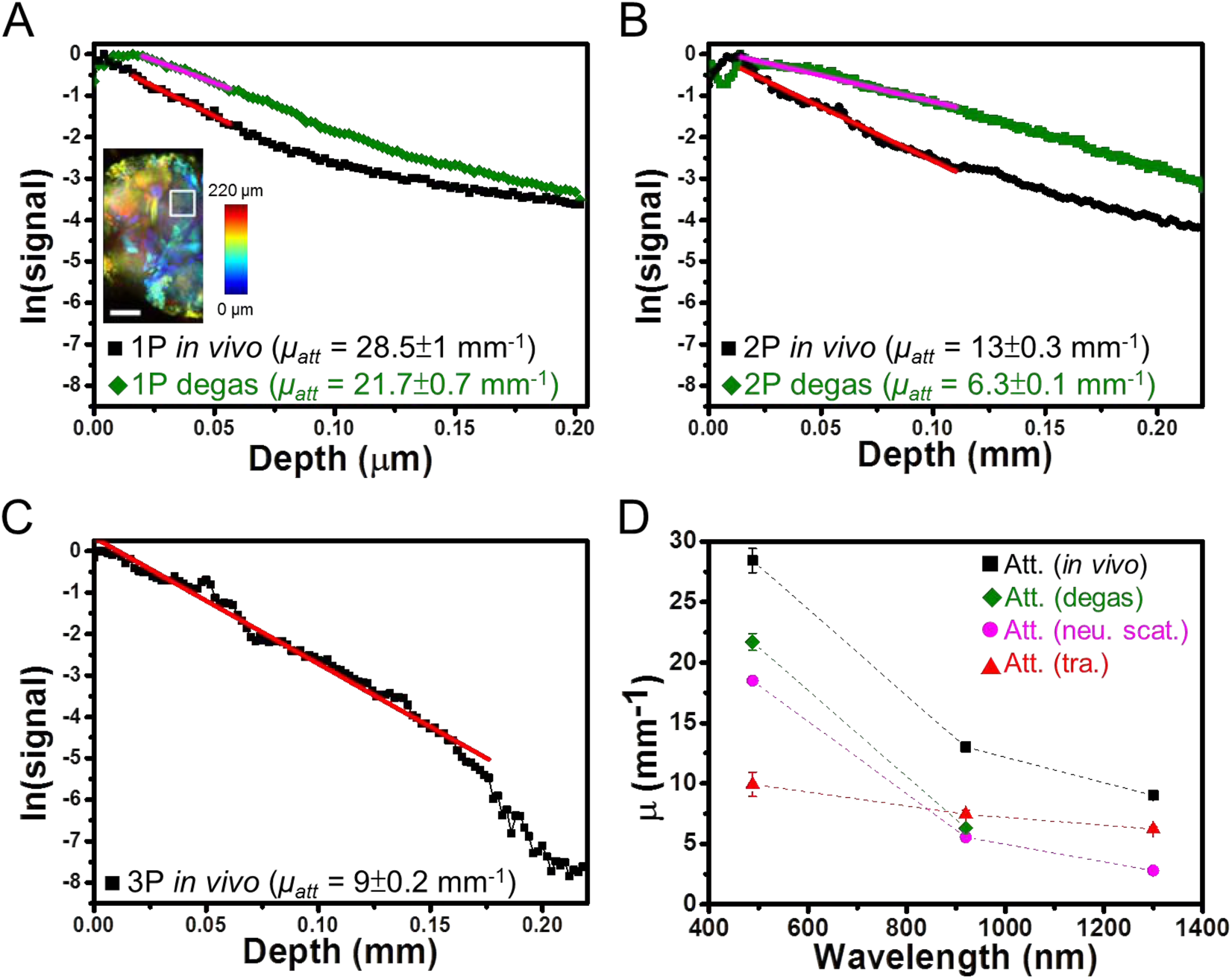
1PF, 2PF, and 3PF signal attenuations in living and degassed brains, and quantitative analysis on contributions from scattering and aberration. (A) and (B) show the semi-logarithmic plot of 1PF and 2PF signal attenuations with depth, inside a living (black) and degassed (green) brains, respectively corresponding to Figure 1A - 1D. (C) 3PF signal attenuation of a living brain in Figure 1E. The inset in (A) shows the area (white box) for signal extraction at different depths. The signals are obtained by averaging the brightest 1 % pixels at different depths inside the white box. In (A) - (C), the corresponding attenuation coefficients (*μ_att_*) are given in the bottom. (D) Quantitative comparison of *μ_att_* in living brains (black squares) and degassed brains (green diamonds) at each wavelength, together with corresponding attenuations from mouse brains, i.e. neuron scattering (pink circles) and tracheae (red triangles). Scale bar in inset of (A): 50 μm.

In Figure 2D, the attenuation coefficients of *in vivo* and degassed brains are plotted in black and green, respectively. It is interesting to note that the attenuations of degassed brains are comparable to that of a mouse brain (pink circles), which is mostly contributed from scattering of neurons (Horton et al., 2013). The slight difference may be due to the effect of residual wax in the tracheae. From the comparison of *in vivo Drosophila* brain (total attenuation, *μ_att_*) and mouse brain (attenuation from neurons, *μ_neu_*), the contribution of trachea *μ_tra_* = *μ_att_* - *μ_neu_* is derived as red triangles in Figure 2D.

The results show that both *μ_neu_* and *μ_tra_* reduce with increasing wavelengths, with respectively λ^−2^ and λ^−05^ dependencies (see Figure supplement 2). Since the attenuation of neuronal tissue *μneu* is dominated by scattering, which includes Rayleigh scattering (λ^−4^ dependence) and Mie scattering (λ^−1^ dependence), the λ^−2^ dependency should be a natural result of the mixture. On the other hand, the λ^−0.5^ dependency suggests that *μ_tra_* may not be governed by scattering, whose wavelength dependency should be in between λ^−4^ and λ^−1^, but by aberration from air-tissue interfaces.

To further confirm that scattering does not play the key role in the attenuation of the trachea-filled brain, we compare the 2PF *μ_att_* of *Drosophila* and mouse in Figure 2D, the latter (~ 5.55 mm^−1^) is less than half of the former (~ 13.5 ± 0.5 mm^−1^). Nevertheless, 2PF imaging in mouse brains typically reaches almost 1 mm, i.e. about five scattering lengths; while in living *Drosophila* brains, only about 100 μm, i.e. less than two attenuation lengths, is achieved. Similar situation occurs with the 3PF modality. The surprisingly limited penetration depth suggests that scattering does not dominate the attenuation, but the trachea-induced aberration dictates, especially in multiphoton imaging cases. The underlying reason of the limited depth is that aberration not only induce excitation intensity loss, but also strongly deteriorates point-spread-function (PSF), especially in the axial direction (Figure supplement 3). As a result, capability of 2PF optical sectioning quickly loses at relatively shallow region, leading to blurred images and considerably limited imaging depth, as manifested in Figure 1C. On the other hand, 3PF imaging modality not only exhibits less aberration (Figure 2D), but also benefits from better optical sectioning (Figure supplement 3), thus provides deeper imaging depth than 2PF. In contrast, the single-photon fluorescence images in Figure 1A (see the arrow) demonstrate loss of image contrast mainly due to scattering.

In Figure 2A and 2B, the trachea-contributed aberration is more efficiently removed by degassing in the 2PF modality (2-fold reduction in *μ_att_*, Figure 2B) than 1PF (~ 30 % reduction, Figure 2A). In addition, from Figure 2D, *μ_tra_* for 2PF and 3PF are larger than *μ_neu_*, especially in the 3PF case. These two observations further suggest that tracheae-contributed aberration is the main factor to impede multiphoton deep-brain imaging in a living *Drosophila*.

As shown in Figure 1, to reaching the milestone of *in vivo* whole-brain imaging in *Drosophila,* 3PF at 1300-nm should be the optimal choice. Here we quantify the depth limit of 3PF by analyzing its signal-to-background ratio (SBR) in Figure 3A and 3b, along with comparison between 2PF and 3PF (see Materials and Methods for SBR calculations). The former reaches unity at ~ 100 μm, since no structure is distinguishable beyond this depth (Video 2), while the SBR of the latter significantly outstands the former. From Figure 3B, the imaging depth of 3PF reaches at least 200 μm. The result once again supports that 3PF is capable to image through a whole *Drosophila* brain.

**Figure 3.**
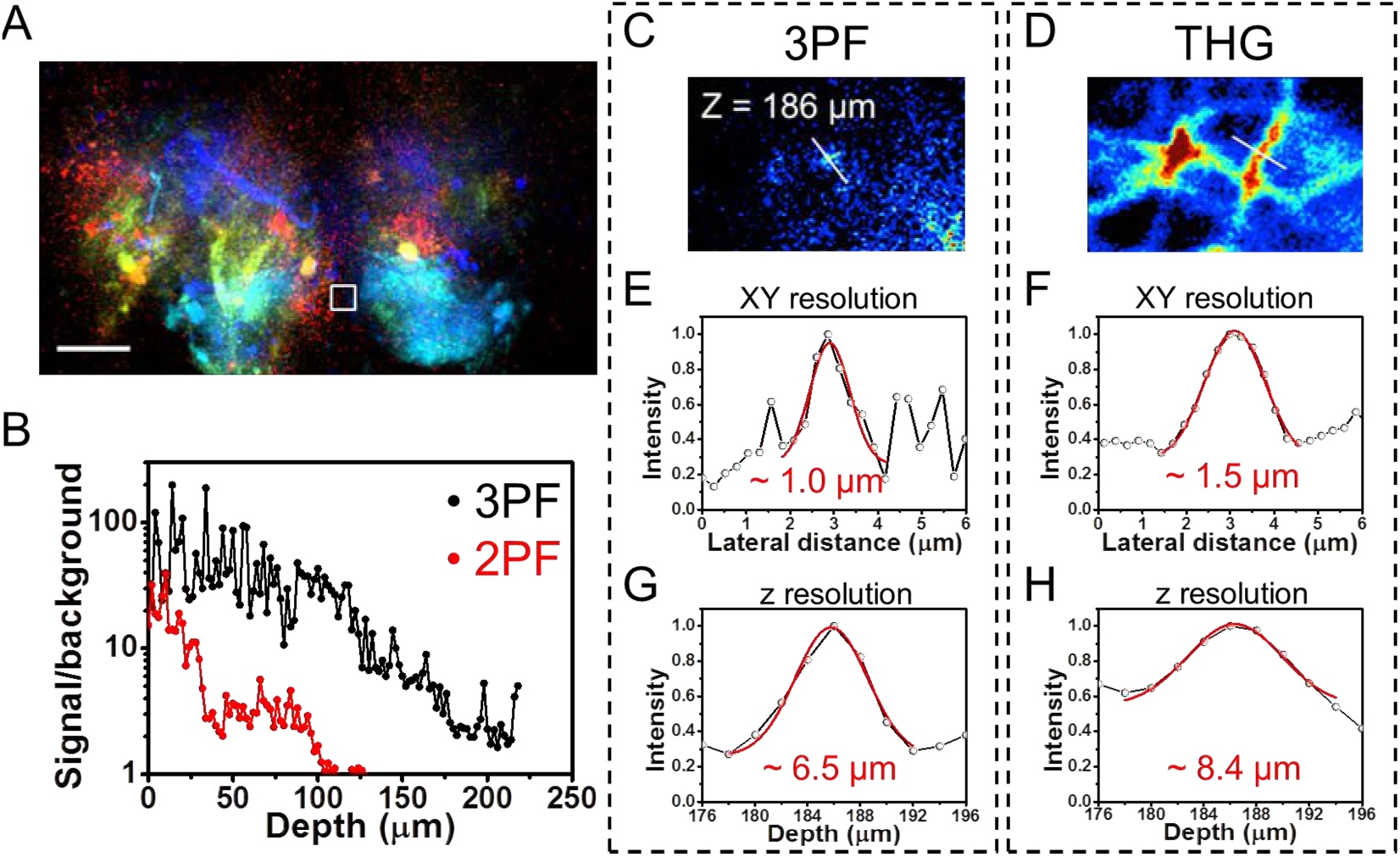
Signal-to-background ratio (SBR) and resolution analysis. (A) Color-coded depth projection image of 3PF throughout the whole-brain, showing different structures at different depths. The white box in the center represents the region of background selection. (B) SBR of 2PF and 3PF signals at each depth. (C) and (D) show the depth images of 3PF and THG at 186 μm, respectively. The lateral resolutions are ~1.0 μm (E) for 3PF and ~ 1.5 μm (F) for THG signals. The axial resolutions are ~ 6.5 μm for 3PF (G) and ~ 8.4 μm for THG (H), respectively. Scale bars: 50 μm in (A), 5 μm in (C).

The spatial resolutions of 3PF and THG in deep-brain region (186 μm depth) are given in Figure 3C - 3H. For 3PF, the minimal full-width-at-half-maximum (FWHM) is 1.0 μm laterally (Figure 3E) and 6.5 μm axially (Figure 3G), which is sufficient to resolve a single cell in the *Drosophila* brain(Tuthill, 2009). For THG, the FWHM is slightly larger, 1.5 μm in lateral (Figure 3F) and 8.4 μm in axial directions (Figure 3H). Considering the width of tracheae, the results should be considered as the upper bound of the THG spatial resolution.

In conclusion, we have, for the first time, characterized the optical properties of the *Drosophila* brain, which is filled with air, with single-photon, two-photon, and three-photon modalities. We found that the main limiting factor that impedes *in vivo* whole-brain single-photon imaging is scattering, but for multiphoton imaging, aberration from tracheae structures plays a more dominant role. The aberration affects not only signal attenuation, but also image visibility. Although degassing enables whole-Drosophila-brain imaging by reducing trachea aberration, the only way to achieve *in vivo* whole-brain imaging with single cell resolution is 3PF at 1300-nm excitation, which exhibits less scattering, aberration, and better optical sectioning. It is possible to combine with AO to further reduce aberration (Tao et al., 2017), thus allowing functional imaging on the scale extending from a single neuron, a complete brain network, toward a whole-animal connectome (Lo & Chiang, 2016).

## Materials and Methods

### Microscope setup

#### One-photon microscope

The one-photon imaging was done on a commercial microscope LSM 780 (Zeiss, Germany). The built-in laser (488-nm) and photomultiplier tube was used to single-photon excitation and signal detection. A water immersion objective was used (Olympus, XLPlan N, 25× NA 1.05) for its high transmission in both visible and IR wavelength ranges. A pinhole with ~ 60 μm diameter was used to achieve optical sectioning. The image formation was done by the controlling software Zen (Zeiss, Germany).

#### Multiphoton microscope

The setup was the same as that in Ouzounov et al (Ouzounov et al., 2017). A home-built laser-scanning microscope that is compatible to long wavelength excitation is constructed. A Ti: sapphire laser at 920-nm with an 80 MHz repetition rate, and an optical parametric amplifier at 1300-nm with a 400 kHz repetition rate, were used as excitation sources of 2PF and 3PF respectively. The same water immersion objective as single-photon microscope was used. The power levels for both lasers after the objective were limited to less than 20 mW for all imaging depths. The fluorescence and THG signals were epi-collected with a dichroic beamsplitter (Semrock, FF705-Di01-25 × 36), and then detected by a GaAsP photomultiplier tube (Hamamatsu, H7422-40) and a bialkali photomultiplier tube (Hamamatsu, R7600-200) in non-descanned configurations to maximize the collection efficiency. A 488-nm dichroic beamsplitter (Semrock, Di02-R488-25×36) was used to split the fluorescence and THG signals, which were further separated by a 520/60 band-pass filter (BPF, transmission at center 520-nm, FWHM 60 nm) for the fluorescence and a 420/40 BPF for the THG. A living *Drosophila* was fixed and placed onto a motorized stage (M-285, Sutter Instrument). A computer running the ScanImage 3.8 under Matlab (MathWorks) was used to synchronize the stage movement and image acquisition.

The signal current from the detectors was converted to voltage, amplified and low-pass filtered by a transimpedance amplifier (Hamamatsu, C9999) and another 1.9 MHz low-pass filter (BLP-1.9+, Minicircuits). Analog-to-digital conversion was performed by a data acquisition card (PCI-6115, National Instruments).

#### Sample preparations

All the sample preparation methods followed the protocol of previous publication on *in vivo Drosophila* brain imaging (H.-H. Lin, Chu, Fu, Dickson, & Chiang, 2013). The samples were adult, female *Drosophila* between 5 and 10 days old. GFP was pan-neuronal expressed by genetic drivers (Gal4-elav.L/CyO × UAS-EGFP). The living *Drosophila* was immobilized in a pipette tip with volume 100 μL after anesthetized by ice bathing. A window was cut into the head by using fine tweezers, after placing a drop of Ca^2+^-free saline on the brain to prevent desiccation, and fat bodies above the brain were removed, under a stereomicroscope. The dissection saline was then replaced with a drop of Ca^2+^-containing saline (108 mM NaCl, 5 mM KCl, 2 mM CaCl2, 8.2 mM MgCl2, 4 mM NaHCO3, 1 mM NaH2PO4, 5 mM trehalose, 10 mM sucrose, and 5 mM HEPES [pH 7.5, 265 mOsm]). No cover glass was placed between the brain and the objective.

To check the optical effect caused by the tracheae structure, degassing the *Drosophila* brain was performed by pumping out the air inside tracheae. The degassing protocol followed the previous publication of *in situ Drosophila* brain imaging (C.-W. Lin et al., 2015). The degassing protocol started from immersing the *Drosophila* in 4 % paraformaldehyde and 2 % triton, expelling air in tracheae by using a vacuum chamber that was depressurized to – 72-mmHg for 2.5 minutes, wait for 1.5 minutes, and then releasing to normal pressure for 2 minutes. The degassing process was completed by repeating the above procedure 4 times. After degassing, the same microsurgery preparation was performed, and observed under the same microscope.

### Signal analyses

#### Attenuation coefficient calculation

This section explains how to derive the attenuation coefficients (*μ_att_*), which is the inverse value of attenuation length (*l_a_*), i.e., *μ_att_* = *l_a_*^−1^, in the results of Figure 2. The calculation of attenuation length inside a biological tissue has been detailed in an earlier work (Horton et al., 2013). The well-known light attenuation equation is:

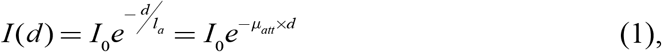

where *I*(*d*) is the excitation intensity at imaging depth of *d,* and *I_0_* is the intensity at tissue surface. With fluorescence excitation processes, the *n*-th order fluorescence intensity, *F*^(*n*)^(*d*), is related to the excitation intensity as:

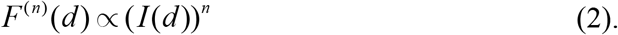

For 1PF, 2PF and 3PF, *n* equals 1, 2 and 3, respectively. Combining eqs. 1 and 2, *F*^(*n*)^(*d*) is obtain in eq. 3,

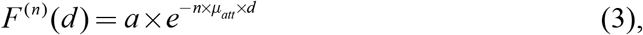

where *a* is a proportional constant. *F*^(*n*)^(*d*) and *d* are determined experimentally, and *μ_att_* can be obtained from their dependencies. More explicitly, by taking the natural logarithmic value of both sides in eq. 3, it becomes,

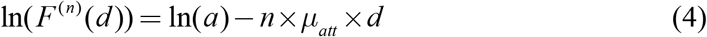

By plotting the dependence of ln(*F*^(*n*)^(*d*)) on *d,* as shown in Figure 2A – 2C and Figure supplement 1B - 1D, the decay slopes of the curves are – *n* × *μ_att_*.

Therefore, *μ_att_* is determined by dividing the inverse value of the slopes over *n*.

To obtain the slopes of the curves in Figure 2A - 2C, the data points for linear regression fitting are selected by the same criteria. They are all selected with the starting point of signal decay to the depth limit of the corresponding imaging modality. For example, the depth range for fitting in Figure 2B starts ~ 20 μm and ends ~ 110 μm. However, the imaging depth limit of 3PF represent the bottom of the brain, instead of the penetration depth limit.

#### Signal-to-background (SBR) calculation

The SBR calculation was used in Figure 3 to determine the imaging depth limitation, and the method was similar to our previous publication (Ouzounov et al., 2017). Briefly, the signal value was averaged from the pixels with top 1 % intensities from the white box in inset of Figure 2A, and the background value was the average of all pixel intensities in the white box of Figure 3A, where no brain tissue was found throughout the whole depth range. With this definition, the signals mostly came from the in-focus regions, and the background arose from out-of-imaging-plane contributions. Both SBR results are normalized to the 2PF value at 100 μm depth, where no distinguishable features are observable.

## Acknowledgements

The three-photon brain imaging cannot be done with the great help from Chris Xu and Tianyu Wang from School of Applied and Engineering Physics in Cornell University. In addition, we appreciate the generous support of transgenic *Drosophila* from Chun Han and Rushaniya Fazliyeva from Institute of Cell and Molecular Biology in Cornell University. This work was financially supported by the Ministry of Science and Technology (MOST), Taiwan, under grant MOST-107-2321-B-002-009-, MOST-105-2628-M-002-010-MY4 and MOST 104-2218-E-007 −022 -MY2, and by the Brain Research Center from The Featured Areas Research Center Program within the framework of the Higher Education Sprout Project by the Ministry of Education (MOE) and MOST in Taiwan. SWC acknowledge the generous support from the Foundation for the Advancement of Outstanding Scholarship.

---

**Figure supplement 1.**
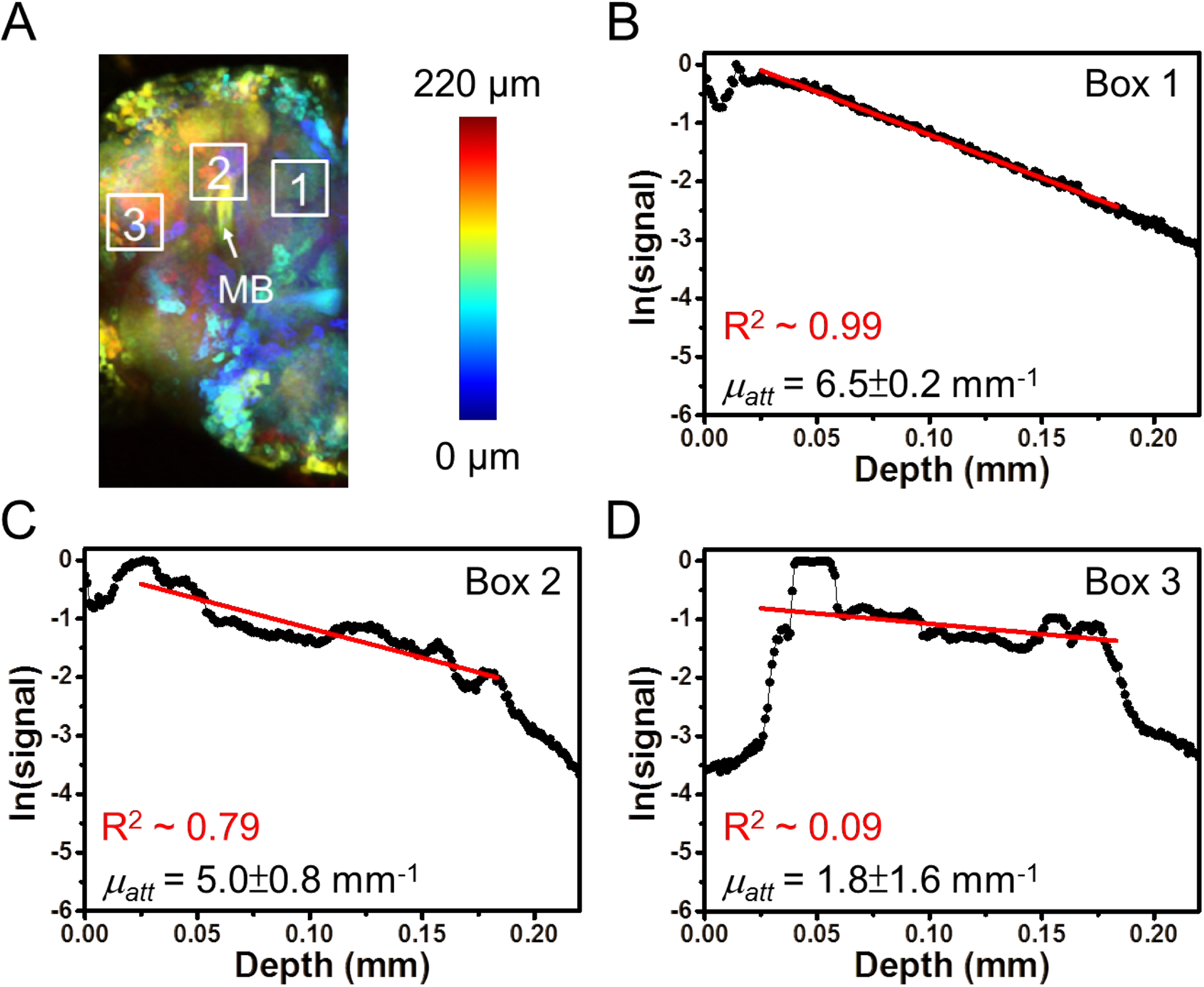

**Figure supp 2.**
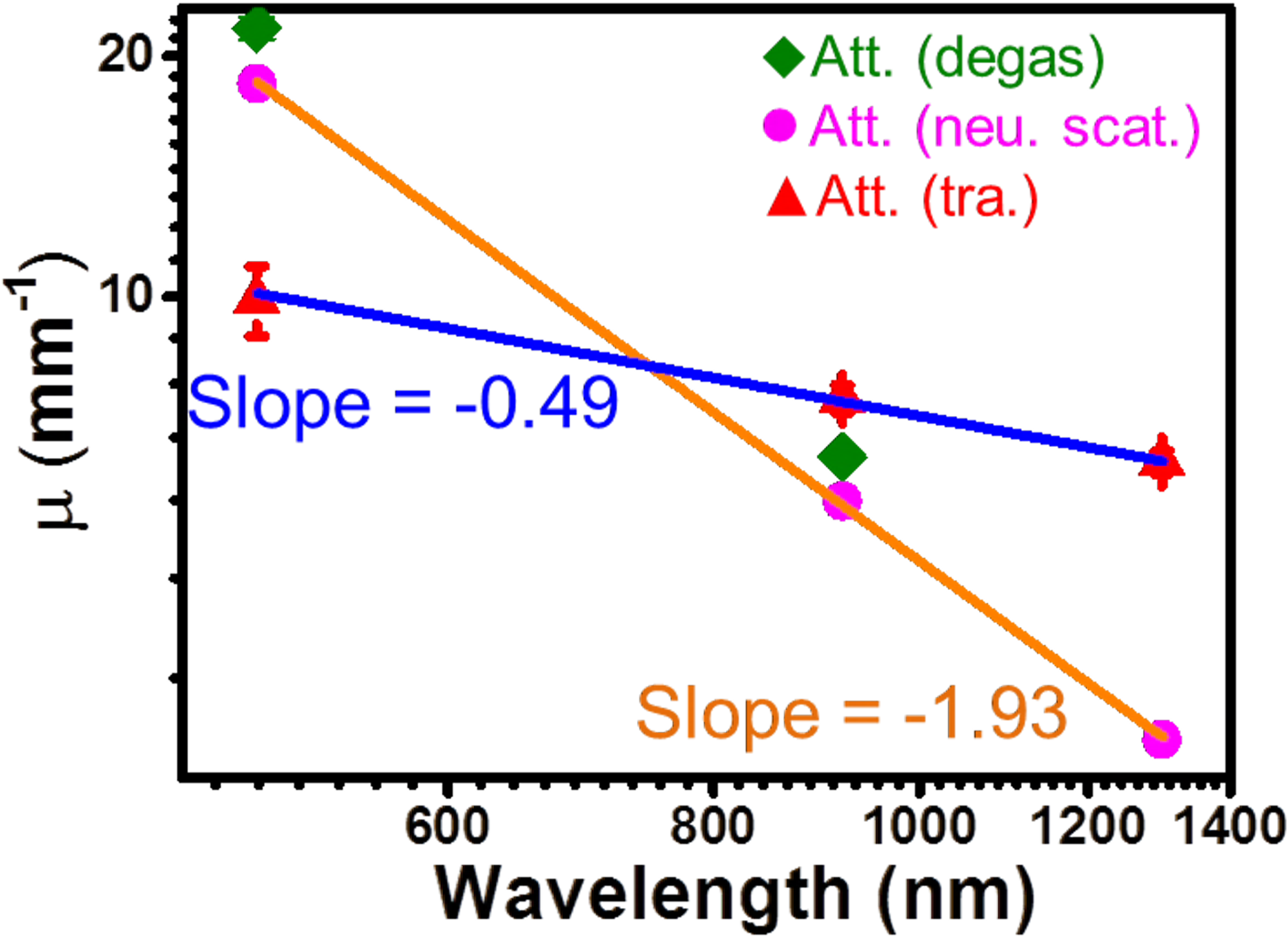

**Figure supplement 3.**
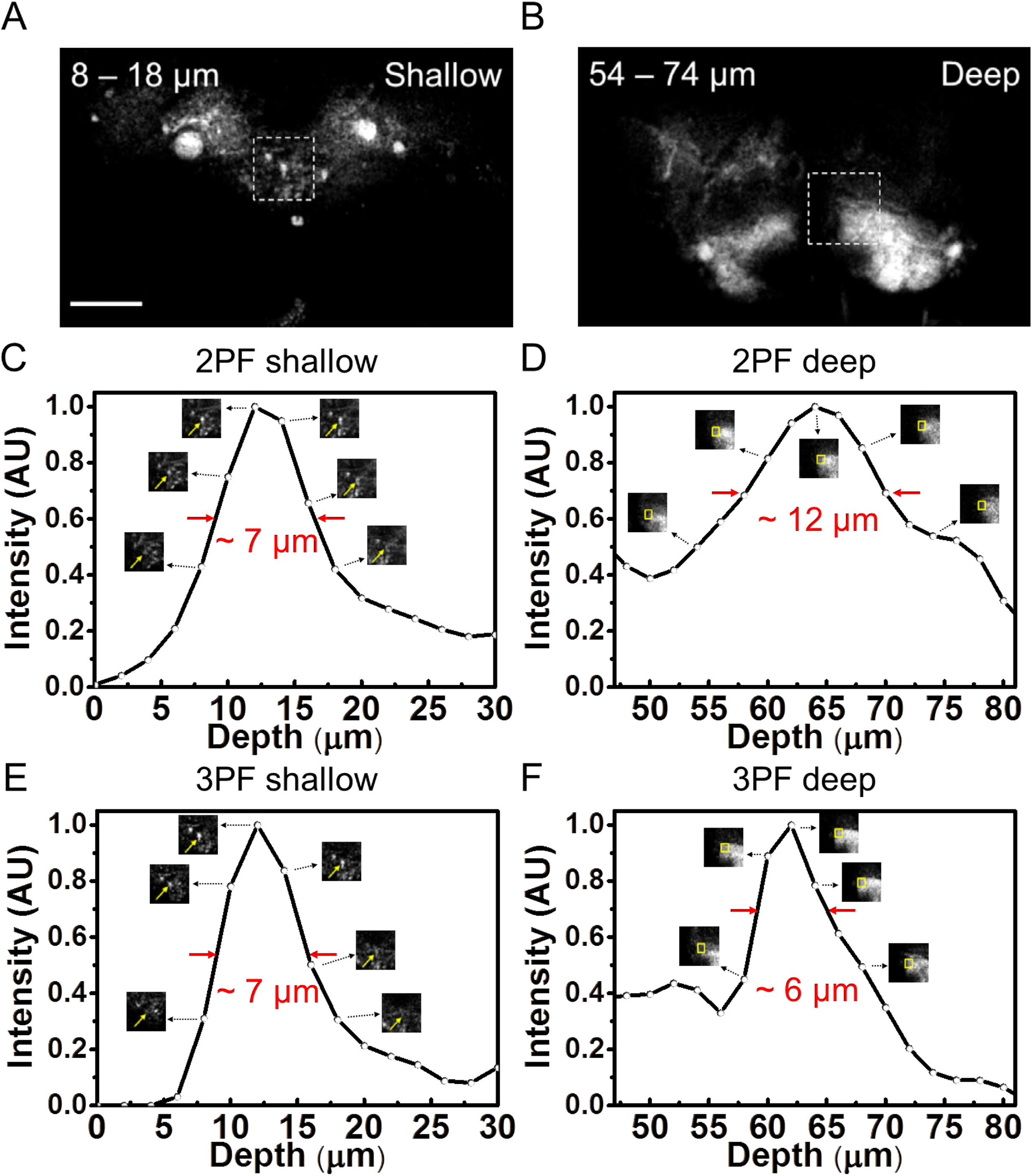

## References

Beitel G. J., & Krasnow M. A. 2000. Genetic control of epithelial tube size in the *Drosophila* tracheal system. Development 127: 3271–3282.

Booth M. J. 2014. Adaptive optical microscopy: the ongoing quest for a perfect image. Light, science & applications 3: e165. DOI: http://doi.org/10.1038/lsa.2014.46

Chiang A.-S., Lin C.-Y., Chuang C.-C., Chang H.-M., Hsieh C.-H., Yeh C.-W., … Hwang J.-K. 2011. Three-dimensional reconstruction of brain-wide wiring networks in *Drosophila* at single-cell resolution. Current Biology 21: 1–11. DOI: https://doi.org/10.1016/j.cub.2010.11.056

Chu S.-W., Chen S.-Y., Tsai T.-H., Liu T.-M., Lin C.-Y., Tsai H.-J., & Sun C.-K. 2003. *In vivo* developmental biology study using noninvasive multi-harmonic generation microscopy. Optics Express 11: 3093–3099. DOI: https://doi.org/10.1364/OE.11.003093

Helmchen F., & Denk W. 2005. Deep tissue two-photon microscopy. Nature Methods 2: 932–940. DOI: https://doi.org/10.1038/nmeth818

Honegger K. S., Campbell R. A. A., & Turner G. C. 2011. Cellular-resolution population imaging reveals robust sparse coding in the *Drosophila*mushroom body. Journal of Neuroscience 31: 11772–11785. DOI: https://doi.org/10.1523/JNEUROSCI.1099-11.2011

Horton N. G., Wang K., Kobat D., Clark C. G., Wise F. W., Schaffer C. B., & Xu C. 2013. *In vivo* three-photon microscopy of subcortical structures within an intact mouse brain. Nature Photonics 7: 205–209. DOI: https://doi.org/10.1038/nphoton.2012.336

Ignell R., Root C. M., Birse R. T., Wang J. W., Nässel D. R., & Winther Å. M. E. 2009. Presynaptic peptidergic modulation of olfactory receptor neurons in *Drosophila*. Proceedings of the National Academy of Sciences 106: 13070–13075. DOI: https://doi.org/10.1073/pnas.0813004106

Kobat D., Horton N. G., & Xu C. 2011. *In vivo* two-photon microscopy to 1.6-mm depth in mouse cortex. Journal of Biomedical Optics 16: 106014. DOI: http://doi.org/10.1117/1.3646209

Lin C.-W., Lin H.-W., Chiu M.-T., Shih Y.-H., Wang T.-Y., Chang H.-M., & Chiang A.-S. 2015. Automated *in situ* brain imaging for mapping the *Drosophila* connectome. Journal of Neurogenetics 29: 157–168. DOI: http://doi.org/10.3109/01677063.2015.1078801

Lin H.-H., Chu L.-A., Fu T.-F., Dickson B. J., & Chiang A.-S. 2013. Parallel neural pathways mediate CO_2_ avoidance responses in *Drosophila*. Science 340: 1338–1341. DOI: http://doi.org/10.1126/science.1236693

Lo C.-C., & Chiang A.-S. 2016. Toward whole-body connectomics. The Journal of Neuroscience 36: 11375–11383. DOI: http://doi.org/10.1523/jneurosci.2930-16.2016

Ouzounov D. G., Wang T., Wang M., Feng D. D., Horton N. G., Cruz-Hernandez J. C., … Xu C. 2017.*In vivo* three-photon imaging of activity of GCaMP6-labeled neurons deep in intact mouse brain. Nature Methods 14: 388–390. DOI: http://doi.org/10.1038/nmeth.4183

Pedrazzani M., Loriette V., Tchenio P., Benrezzak S., Nutarelli D., & Fragola A. 2016. Sensorless adaptive optics implementation in widefield optical sectioning microscopy inside*in vivo Drosophila* brain. Journal of Biomedical Optics 21: 036006. DOI: http://doi.org/10.1117/1.JBO.21.3.036006

Root C. M., Semmelhack J. L., Wong A. M., Flores J., & Wang J. W. 2007. Propagation of olfactory information in *Drosophila*. Proceedings of the National Academy of Sciences 104: 11826–11831. DOI: https://doi.org/10.1073/pnas.0704523104

Ruta V., Datta S. R., Vasconcelos M. L., Freeland J., Looger L. L., & Axel R. 2010. A dimorphic pheromone circuit in *Drosophila* from sensory input to descending output. Nature 468: 686–690. DOI: https://doi.org/10.1038/nature09554

Tang J., Germain R. N., & Cui M. 2012. Superpenetration optical microscopy by iterative multiphoton adaptive compensation technique. Proceedings of the National Academy of Sciences 109: 8434–8439. DOI: http://doi.org/10.1073/pnas.1119590109

Tao X., Lin H.-H., Lam T., Rodriguez R., Wang J. W., & Kubby J. 2017. Transcutical imaging with cellular and subcellular resolution. Biomedical Optics Express 8: 1277–1289. DOI: http://doi.org/10.1364/BOE.8.001277

Theer P., Hasan M. T., & Denk W. 2003. Two-photon imaging to a depth of 1000 μm in living brains by use of a Ti: Al_2_O_3_ regenerative amplifier. Optics Letters 28: 1022–1024. DOI: https://doi.org/10.1364/OL.28.001022

Tuthill J. C. 2009. Lessons from a compartmental model of a *Drosophila* neuron. Journal of Neuroscience 29: 12033–12034. DOI: https://doi.org/10.1523/JNEUROSCI.3348-09.2009

Wang C., Liu R., Milkie D. E., Sun W., Tan Z., Kerlin A., … Ji N. 2014. Multiplexed aberration measurement for deep tissue imaging *in vivo*. Nature Methods 11: 1037–1040. DOI: http://doi.org/10.1038/nmeth.3068

Wang J. W., Wong A. M., Flores J., Vosshall L. B., & Axel R. 2003. Two-photon calcium imaging reveals an odor-evoked map of activity in the fly brain. Cell 112: 271–282. DOI: https://doi.org/10.1016/S0092-8674(03)00004-7

Wang Y. L., Guo H. F., Pologruto T. A., Hannan F., Hakker I., Svoboda K., & Zhong Y. 2004. Stereotyped odor-evoked activity in the mushroom body of *Drosophila* revealed by green fluorescent protein-based Ca^2+^ imaging. Journal of Neuroscience 24: 6507–6514. DOI: https://doi.org/10.1523/JNEUROSCI.3727-03.2004

Wei L., Chen Z., & Min W. 2012. Stimulated emission reduced fluorescence microscopy: a concept for extending the fundamental depth limit of two-photon fluorescence imaging. Biomedical Optics Express 3: 1465–1475. DOI: http://doi.org/10.1364/BOE.3.001465

